# Transcriptome and DNA methylome dynamics reveal differential characteristics of inflorescence development between two ecotypes in *Panicum hallii*

**DOI:** 10.1101/2022.02.28.482119

**Authors:** Xiaoyu Weng, Haili Song, Avinash Sreedasyam, Taslima Haque, Li Zhang, Cindy Chen, Yuko Yoshinaga, Melissa Williams, Ronan C. O’Malley, Jane Grimwood, Jeremy Schmutz, Thomas E. Juenger

**Affiliations:** Department of Integrative Biology, University of Texas at Austin, Austin, TX, USA; HudsonAlpha Institute for Biotechnology, Huntsville, AL, USA; US Department of Energy Joint Genome Institute, Lawrence Berkeley National Laboratory, Berkeley, CA, USA

## Abstract

The morphological diversity of the inflorescence determines flower and seed production, which is critical for plant adaptation and fitness. Cytosine methylation is an epigenetic mark that contributes to gene expression regulation during inflorescence development. *Panicum hallii* is a wild perennial grass in the subfamily Panicoideae that has been developed as a model to study perennial grass biology and adaptive evolution. Highly divergent inflorescences have evolved between the two major ecotypes in *P. hallii*, the upland ecotype with compact inflorescence and large seed and the lowland ecotype with an open inflorescence and small seed. Here we performed a comparative transcriptome and DNA methylome analysis across different stages of inflorescence between these two divergent ecotypes of *P. hallii*. Global transcriptome analysis identified differentially expressed genes (DEGs) involved in panicle divergence, stage-specific expression, and co-expression modules underlying inflorescence development. Comparing DNA methylome profiles revealed a remarkable level of differential DNA methylation associated with the evolution of the *P. hallii* inflorescence. We found that most differentially methylated regions (DMRs) occurred within the flanking regulatory regions of genes, especially the promoter. Integrative analysis of DEGs and DMRs characterized the global features of DMR-associated DEGs in the divergence of *P. hallii* inflorescence, which includes homologs of important inflorescence and seed developmental genes that have been previously identified in domesticated crops. Evolutionary analysis measured by *Ka*/*Ks* ratio suggested that most DMR-associated DEGs are under relatively strong purifying selection. This study provides insights into the transcriptome and epigenetic landscape of inflorescence divergence in *P. hallii* and a novel genomic resource for perennial grass biology.

**One sentence summary:** A comparative transcriptome and DNA methylome analysis of inflorescence between upland and lowland ecotypes reveal gene expression and DNA methylation variation underlying inflorescence divergence in *Panicum hallii*.

## Introduction

Flowering plants have evolved diverse inflorescences architecture which determines the spatial arrangement of inflorescence branching and flower production and dictates the production of seed set (Harder and Prusinkiewicz, 2013). The extensive diversity of inflorescence architecture is shaped by a combination of genetic, epigenetic, and environmental factors, with critical economic significance in agricultural and also profound ecological implications in wild species (Barazesh and McSteen, 2008; Teo et al., 2014; Tu et al., 2019). Recently, progress has been made in understanding the area of natural genetic architecture underlying inflorescence development, largely focusing on the model plants *Arabidopsis* and several important crops, including rice, maize, wheat and Setaria (Kellogg et al., 2013; Zhang and Yuan, 2014; Li et al., 2021). This accumulated knowledge provides an opportunity to better understand the role of inflorescence diversity in the adaptive evolution of wild plants.

DNA methylation is a heritable epigenetic modification that contributes to gene regulation and genome structure and integrity (Chan et al., 2005; Law and Jacobsen, 2010; Zhang et al., 2018). In land plants, DNA methylation occurs at the cytosine bases with three sequence contexts (CG, CHG and CHH, where H represents A, T or C). Genome-scale DNA methylation analyses show extensive variation among different plant species in all three DNA methylation contexts, with the predominant form being CG methylation compared with CHG and CHH methylation (Niederhuth et al., 2016). The classic model assumes that the addition of DNA methylation in the promoters of genes typically represses gene expression by recruiting repressor proteins (Tate and Bird, 1993). Recently, a growing body of research has revealed that gene body methylation can be positively correlated with gene expression and may shape important features of plant genome evolution (Bewick and Schmitz, 2017). DNA methylation plays an essential role in a wide range of growth and development events, especially in the developmental complexity of inflorescence architecture (Zhang et al., 2018; Tu et al., 2019). This perspective is supported by evidence that most loss-of-function mutations of genes involving in DNA methylation establishment and maintenance show abnormal inflorescence morphology (Moritoh et al., 2012; Fernandez-Nohales et al., 2014; Liao et al., 2019). Additionally, epigenetic alleles involving DNA methylation variation have been identified in the key regulators of inflorescence development. Among them, the *ipa1-2D* allele was associated with a reduction of DNA methylation at the promoter of *IDEAL PLANT ARCHITECTURE1* (*IPA1*), which encodes the protein SPL14 (SQUAMOSA-promoter binding protein-like 14) controlling rice inflorescence branching, thus alleviating the epigenetic suppression of *IPA1* transcription (Zhang et al., 2017). In addition, the expression of *OsLG1* is down-regulated by DNA methylation in its promoter region, which results in dense panicles in domesticated rice cultivars (Zhu et al., 2013). Further, differential CG methylation at the distal regulatory region of *TEOSINTE BRANCHED1* (*TB1*), which encodes a TCP (Teosinte Branched1/Cycloidea/Proliferating cell factor) transcription factor controlling the axillary structures during the maize domestication, acts like a *cis*-acting element in teosinte-maize evolution (Xu et al., 2020). These findings suggest that DNA methylation may be of crucial importance in the evolution of the structure and organization of the inflorescence.

High-throughput sequencing techniques have been used extensively for genome-wide profiling of gene expression and DNA methylation to study a variety of developmental processes (Yang et al., 2015; Huang et al., 2019; Rajkumar et al., 2020; Zhou et al., 2020). As the key determinant of productivity, the inflorescence of many crop species has been studied with detailed developmental stage-specific transcriptome profiling (Furutani et al., 2006; Wang et al., 2010; Eveland et al., 2014; Harrop et al., 2016; Feng et al., 2017; Zhu et al., 2018). For example, stage- and meristem-specific gene expression profiles have provided a genome-wide view of regulatory networks controlling young panicle development in rice (Furutani et al., 2006; Wang et al., 2010; Harrop et al., 2016) and wheat (Feng et al., 2017). Moreover, whole-genome analysis of DNA methylation has found epigenetic mechanisms that coordinate gene structure and expression during inflorescence development (Li et al., 2012; Parvathaneni et al., 2020; Sun et al., 2020). For instance, single-base resolution methylome studies have assessed the functional significance of epigenetic differentiation of young panicle between wild and cultivated rice (Li et al., 2012). Genome-wide DNA methylation profiling integrated with other multi-omics analysis has revealed the role of chromatin interactions that coordinate *trans* and *cis* regulation of differential expression between two separate types of inflorescence (ear and tassel) in maize (Sun et al., 2020). These advances provide not only a deep understanding of the relationship between complex gene regulatory networks and epigenetic modifications but also help to identify the potential candidates controlling inflorescence morphology and grain yield.

*Panicum hallii* (*P. hallii*) is a native perennial C4 grass with a distribution in southwestern regions of North America (Lowry et al., 2015). Due to a close evolutionary relationship to the polyploid biofuel crop switchgrass (*Panicum virgatum*), *P. hallii* has been developed as a complementary diploid model system (Lovell et al., 2018). *P. hallii* is found in a wide range of soil and ecological conditions, spanning from xeric inland regions to mesic coastal areas (Gould et al., 2018; Palacio-Mejia et al., 2021). *P. hallii* populations have diverged into two major ecotypes (or varieties), *P. hallii var. hallii* (hereafter *var. hallii*) and *P. hallii var. filipes* (hereafter *var. filipes*) (Lowry et al., 2015; Lovell et al., 2018). Similar to other upland plants, the widespread *var. hallii* is typically found in drier habitats with shallow and rocky soils (Palacio-Mejia et al., 2021). In contrast, the more geographically restricted *var. filipes* commonly grows in Gulf coast areas in clay soils and mesic depressions (Palacio-Mejia et al., 2021). Whole genome sequencing and assemblies suggest that *var hallii* and *var filipes* shared a common ancestor ∼1.08 million years ago (Lovell et al., 2018). Although there is some evidence of hybridization between these ecotypes, it is rare in nature and they exhibit considerable population structure and genomic and phenotypic divergence, including notable differences in flowering time, plant size and inflorescence architecture (Palacio-Mejia et al., 2021). In general, *var. hallii* flowers earlier than *var. filipes* and is distinguished from the latter by its compact spike-like inflorescence and larger seed. Recently, genetic resources derived from the crossing of *var. hallii* with *var. filipes*have been developed for studying the genetic basis of ecotype-differentiating traits (e.g. flowering time, flower number, seed mass, etc.) and shoot-root resource acquisition traits (Lowry et al., 2015; Khasanova et al., 2019). Razzaque and Juenger (2021, in review) demonstrated local adaptation for *P. hallii* ecotypes in terms of recruitment at the seedling stage at coastal and upland sites, likely resulting from differences in seed mass and dormancy characteristics. Several transcriptome studies have been undertaken with the goal of understanding how *P. hallii* responds to various environmental cues (Lovell et al., 2016; Weng et al., 2019). Nevertheless, gene expression divergence associated with the evolution of ecotype-specific morphology and its relationship with the global patterns of DNA methylation variation remain poorly understood in *P. hallii*.

In this study, we performed a comparative transcriptome and DNA methylome analysis at different stages of inflorescence development contrasting the two ecotypes of *P. hallii* using high-throughput RNA sequencing and whole-genome bisulfite sequencing. Global analysis of transcriptome data identified the patterns of divergence in gene expression between the two types of *P. hallii* inflorescences over development. Similarly, comparing whole-genome DNA methylation profiles allowed a characterization of DNA methylation divergence during the evolution and development of *P. hallii* inflorescence. An integrated analysis of DNA methylation and gene expression highlight important candidate genes that might determine the phenotypic diversity of inflorescence branching architecture and seed size in *P. hallii*. Together, this study provides new insights into transcriptome and epigenetic landscape of inflorescence divergence in *P. hallii*.

## Results

### Distinct phenotypes of inflorescence and seed between two *P. hallii* ecotypes

To provide tools for studying *P. hallii* evolutionary genomics, we have developed reference genomes spanning the wide ecotypic divergence observed in *P. hallii* (Lovell et al., 2018). Our genome assemblies have been derived from two accessions, *P. hallii var. hallii* (HAL2) and *P. hallii var. filipes* (FIL2), that are representative of the upland and lowland ecotypes in *P. hallii* (Lovell et al., 2018). In this study, we investigate inflorescence development between HAL2 and FIL2. As shown in Figure 1A, *P. hallii* has a panicle-type inflorescence with many branches for spikelet development and seed set. The inflorescence of HAL2 exhibits remarkably different branching patterns compared with that in FIL2, mainly in the reduction of both primary and secondary branch numbers (Figure 1A and 1C) (almost spike-like), its compact rather than open structure, and the reduction in the length of spikelet pedicel. This divergent architecture results in a significant decrease in spikelet numbers in HAL2 (Figure 1A and 1C). Further, we observed significantly enlarged seed size in HAL2 relative to FIL2, as measured by hundred-seed-weight (Figure 1B and 1C). We hypothesize that divergence in inflorescence architecture in *P. hallii* may be associated with a trade-off between seed size and number in *P. hallii* as has been observed in many domesticated grasses (Sadras, 2007).

**Figure 1.**
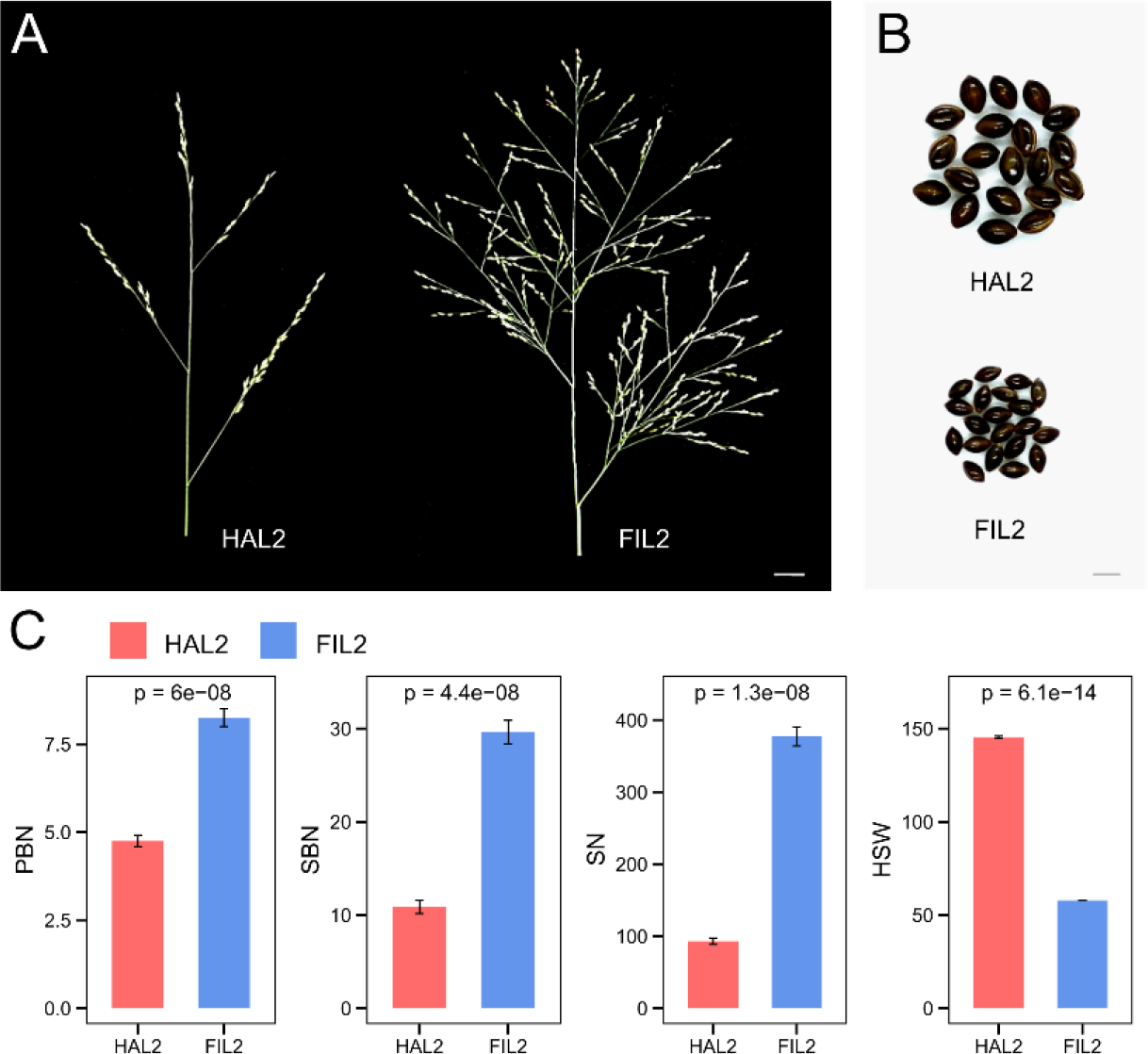
Morphological differences between two representatives of the upland (HAL2) and lowland (FIL2) ecotypes in P. hallii. Representative image of the inflorescence (A) and seed (B) morphology of HAL2 and FIL2 (Scale bar in A, 1 cm; scale bar in B, 1 mm). (C) Primary branch number (PBN), secondary branch number (SBN), spikelet numbers (SN), and hundred-seed-weight (HSW) in HAL2 and FIL2 plants (n = 8).

### Genome-wide analysis of gene expression divergence between two *P. hallii* inflorescences

To explore the molecular mechanisms of inflorescence divergence, four stages of inflorescence tissues from biological replicates of HAL2 and FIL2 were collected for RNA-seq analysis (2 genotypes x 4 developmental stages x 3 biological replicates = 24 libraries). The different stages of inflorescence development were designated as D1-D4 and represented major events (Figure 2A), including the transition of vegetative shoot apical meristem to an inflorescence meristem at the earliest stages, the differentiation of branch and spikelet meristems, and ultimately the generation of the gametes and flower organs at the latest stage. After filtering low expression genes, 19,332 one-to-one orthologous genes were detected in the dataset for downstream analysis. There were strong correlations among the biological replicates (*r* > 0.97), supporting the high quality and reproducibility of the entire dataset. Principal component analysis of expressed genes revealed a strong global structure along the development gradient and related to genotype divergence (Figure 2B). The first component explained 57% of the expression variance and clearly distinguished the stages across the developmental gradient, while the second component explained 35% of the expression variance and mainly discriminated between samples from HAL2 and FIL2 (Figure 2B). The first two components explained the vast majority of variance (92%), suggesting the dominance of development and genotype effects in the entire dataset.

**Figure 2.**
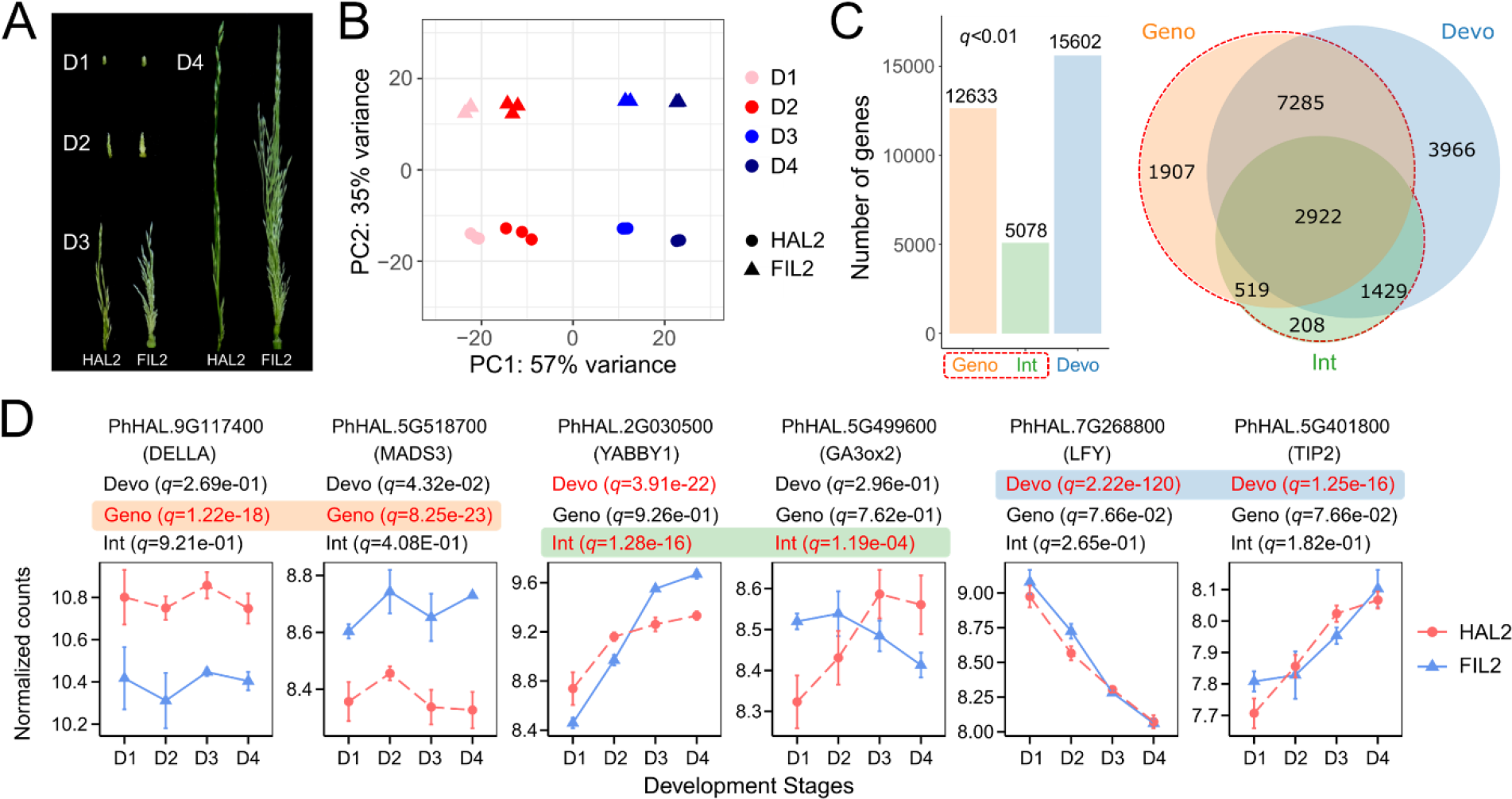
Global transcriptome analysis of gene expression divergence between HAL2 and FIL2 inflorescence. (A) Four stages of inflorescence tissues from HAL2 and FIL2 were collected for RNA-seq. Different stages were designated as D1-D4 and represented major events occurring within young inflorescence development. (B) Principal component analysis of the RNA-seq data for the 24 inflorescence samples showing the developmental signatures and genotype effects. (C) Bar plot and venn diagrams depict genes that are differentially regulated with genotype effects (Geno) development effects (Devo), and genotype-by-development interaction effects (Int) in factorial linear modeling (q-value < 0.01). (D) Expression patterns of development marker genes with significant genotype, development, and genotype-by-development interaction effects. The x axis represents four developmental stages, while the y axis represents normalized counts using variance stabilizing transformation in DEseq2. In all panels, the points and error bars are the average values and SE, respectively, based on normalized counts of three RNA-seq replicates. q-value for each model effect is shown on the top of expression pattern plots.

To investigate the underlying mechanisms of gene expression divergence, we fit linear models on gene counts to test the effects of genotype, development, and genotype × development interaction on gene expression across the entire transcriptome. Genes with significant genotype and/or interaction effects were considered evidence of expression divergence between the ecotypes. In this framework, we identify 14,270 genes (73.8%) with expression divergence between HAL2 and FIL2 inflorescence (Figure 2C, Supplemental Table S1). Among them, 12,633 genes (65.3%) exhibited significant genotype effects (*qgeno* < 0.01) in that they display either genotype and/or interactive effects (Figure 2C, Supplemental Table S1). We only detected 1,907 genes (9.9%) with strictly additive genotype effects, which are genes with consistent difference between HAL2 and FIL2 regardless of developmental stages (Figure 2C, Supplemental Table S1). For examples, the homolog of DELLA protein *SLR1* (PhHAL.9G117400) (*qgeno* = 1.22e-18) (Figure 2D), which regulates panicle architecture via the GA-KNOX signaling pathway in rice (Su et al., 2021), has higher expression in HAL2 at all four sampling stages. In contrast, the homolog of *OsMADS3* (PhHAL.5G518700) (*qgeno* = 8.25e-23) (Figure 2D), which affects maintenance and determinacy of the flower meristem in rice (Li et al., 2011), has higher expression in FIL2 at all four sampling stages. Meanwhile, we detected 5,078 genes (26.3%) with significant interactions between development and genotype (*qint* < 0.01) (Figure 2C, Supplemental Table S1). Here, the magnitude or direction of expression divergence between HAL2 and FIL2 depended on the specific stage of development. Among them, we observed the homologs of rice *YABBY1* (PhHAL.2G030500) (*qint* = 1.28e-16) and *GA3ox2* (PhHAL.5G499600) (*qint* = 1.19e-04) (Figure 2D). It was known that *YABBY1* binds to the promoter of *GA3ox2* in the feedback regulation of GA biosynthesis (Dai et al., 2007). Additionally, we identified 15,602 genes (80.7%) with significant expression level changes across the developmental gradient (*qdevo* < 0.01) (Figure 2C, Supplemental Table S1).

Among them, only 3,966 genes (20.5%) exhibited strictly additive developmental effects without genotype influences (genotype and/or interaction effects) (Figure 2C, Supplemental Table S1). Marker genes controlling inflorescence and floral development were found in this category. For examples, the expression of *LFY* homolog (PhHAL.7G268800) (*qdevo* = 2.22e-120) (Figure 2D), known to regulate meristem initiation at early stages of inflorescence development (Rao et al., 2008), decreased gradually from early to late stages regardless of genotypes. The homolog of *TIP2* (PhHAL.5G401800) (*qdevo* = 1.25e-16) (Figure 2D), known to have crucial roles in anther and pollen development (Fu et al., 2014), showed similar expression patterns between HAL2 and FIL2 with higher expression at late stages than early stages. These results provided a global overview of gene expression divergence and suggests that heterochronic changes in the timing of gene expression may play a critical role in the divergence of HAL2 and FIL2 inflorescence.

### Stage-specific and divergence patterns of gene expression during inflorescence development

To dissect dynamic signals across developmental gradients, we also completed stage-by-stage contrasts of the 14,270 genes with expression divergence between HAL2 and FIL2 inflorescence. We detected 7,843 genes that are differentially expressed under stringent criteria (*q* < 0.01, fold change > 1.5) when contrasting the two genotypes in at least one developmental stage (Figure 3, Supplemental Table S2). The vast majority of these genes (7719) showed expression patterns with consistently greater expression in one of the genotypes (Figure 3); for convenience we call these HAL2 or FIL2 predominant expression patterns. We did not observe a directional bias in the pattern of predominant genes, since 3,963 genes showed HAL2 predominant patterns and 3,756 genes exhibited FIL2 predominant patterns (Figure 3, Supplemental Table S2). We only found 124 genes with rank changing patterns of relative repression or induction changes between genotypes at different development stages (Figure 3, Supplemental Table S2). Further analyses of the stage-specific patterns revealed that 1,075 of HAL2 and 1,050 of FIL2 predominant genes were consistently differentially expressed across all four stages (Figure 3, Supplemental Table S2). We found that 1,350 HAL2 (226 at D1, 160 at D2, 250 at D3, 714 at D4) and 1,298 FIL2 (284 at D1, 166 at D2, 177 at D3, 671 at D4) predominant genes were highly stage-specific and occurred in only one of the four stages (Figure 3, Supplemental Table S2). These analyses suggested that the genotype predominant patterns may be important drivers of the gene expression divergence of inflorescence development between HAL2 and FIL2.

**Figure 3.**
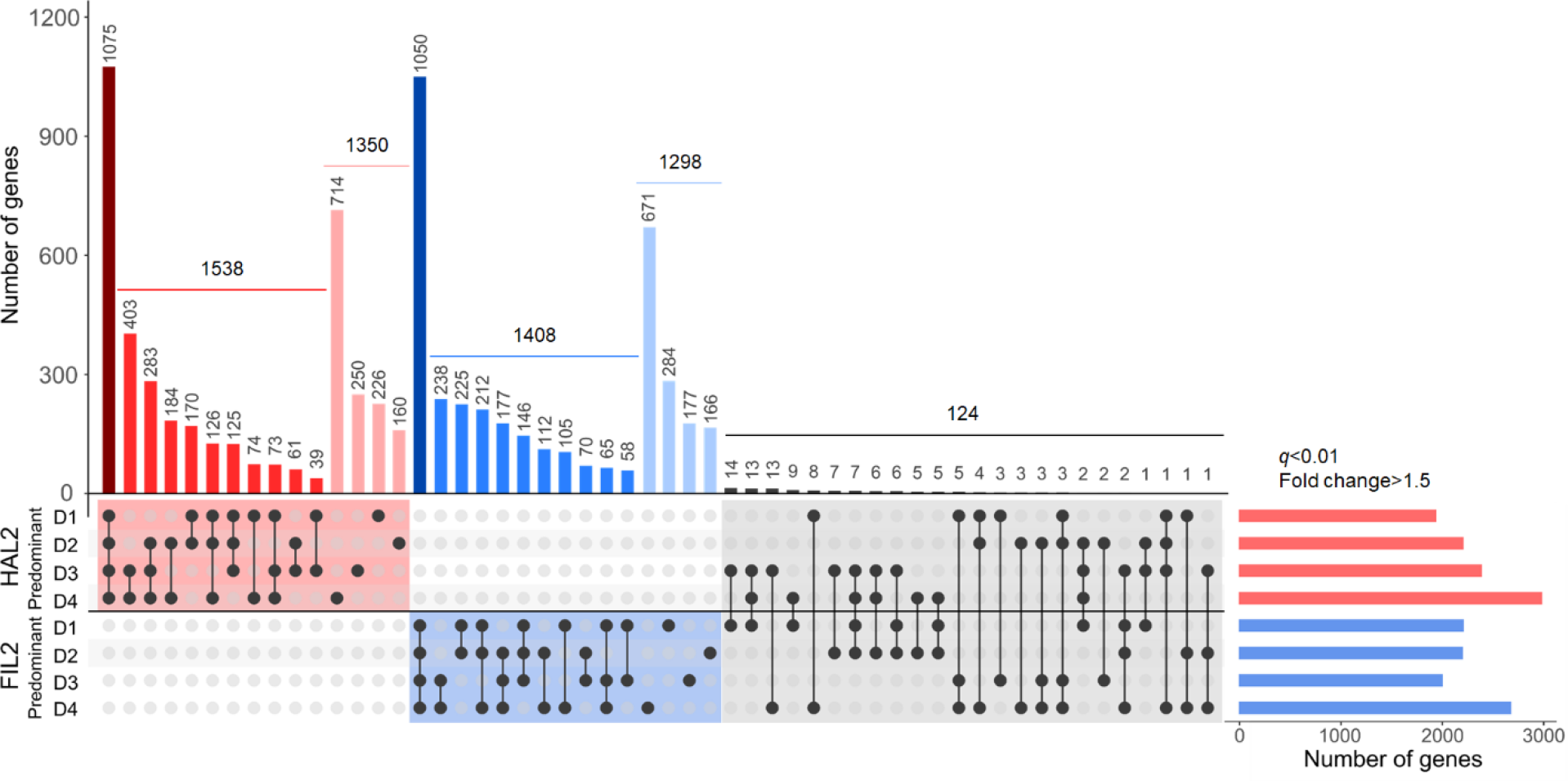
Stage-specific of differentially expressed genes during inflorescence development. Quantification of stage-specific expression of genes in HAL2 predominant (red bar on the top), FIL2 predominant (blue bar on the top), and rank changing (grey bar on the top) patterns. The total numbers of genes that showed higher expression at HAL2 (red bar on the right) or FIL2 (blue bar on the right) at each development stage are represented by horizontal bars on the right of the figure. The numbers of genes showing developmental-specific expression patterns in one or more of sampling stages are shown in black vertical bars of the figure. Black dots at the bottom of each vertical bar indicate the developmental-specific expression identified at each sampling stage. The lined dots indicate two or more sampling stages showing differential expression between two genotypes.

We applied the fuzzy *c*-means algorithm to cluster the 7,843 genes with strong expression divergence (*q* < 0.01, fold change > 1.5) between HAL2 and FIL2 (Kumar and M, 2007). We determine the number of cluster cores (*c*) using the minimum centroid distance with *c* values between 2 to 20, leading to the detection of 10 core clusters representing the divergence pattern of gene expression across development (Supplemental Figure S1). The number of genes in each cluster is presented in Figure 4B and Supplemental Table S2, and ranges from 588 to 1133. Genes from different clusters exhibited opposite dynamic expression trends from early to late stages. Among these, clusters 1, 2, and 9 represented genes that have the tendency to decrease across developmental gradients (Figure 4). We detected a *OsDRM2* homolog (PhHAL.9G458000) and three *SPL* homologs in cluster 1, and eight *MADS-box* homologs in cluster 2 (Supplemental Figure S2). *DRM2* acts as the major DNA methyltransferase in the RNA-directed DNA methylation (RdDM) pathway in plants (Moritoh et al., 2012), *SPL* genes often define the reproductive branching architecture of panicles and therefore flower and seed number (Wang et al., 2015), and *MADS-box* genes play a crucial role from meristems initiation to floret organogenesis (Liu et al., 2013). By contrast, genes in clusters 3, 4, 5, and 8 had expression that gradually increased with maturation of the inflorescence, and genes in clusters 7 and 10 had a sharp increase in expression at the late stages (Figure 4). Numerous hormone-related genes involved in the auxin, gibberellin, and cytokinin metabolism and signaling pathways fell into clusters 3, 4, 5, and 8 (Supplemental Figure S2). The dynamic interactions among these hormones play a fundamental role in the regulation of grass inflorescence architecture (Barazesh and McSteen, 2008). The homolog of *OsDCL2a* (PhHAL.9G171400), a gene that is highly expressed in the egg cell in both monocot and dicot plants (Takanashi et al., 2011), was observed in cluster 7 (Supplemental Figure S2). *DCL2* genes are required for the biosynthesis of small RNA and involved in the RdDM pathway in plants (Yang et al., 2016). Unlike other clusters with increased or decreased expression trends, genes in cluster 6 exhibited relatively stable expression across developmental stages. We noticed that argonaute proteins, such as the homolog of *OsAGO18* (PhHAL.2G357700), and RNA-dependent RNA polymerase, such as the homolog of *OsRDR1* (PhHAL.1G380000), were identified in this cluster (Supplemental Figure S2). These genes play unique roles in small RNA recruitment and biogenesis, and globally alter DNA methylation through RdDM pathway (Kapoor et al., 2008; Wang et al., 2016).

**Figure 4.**
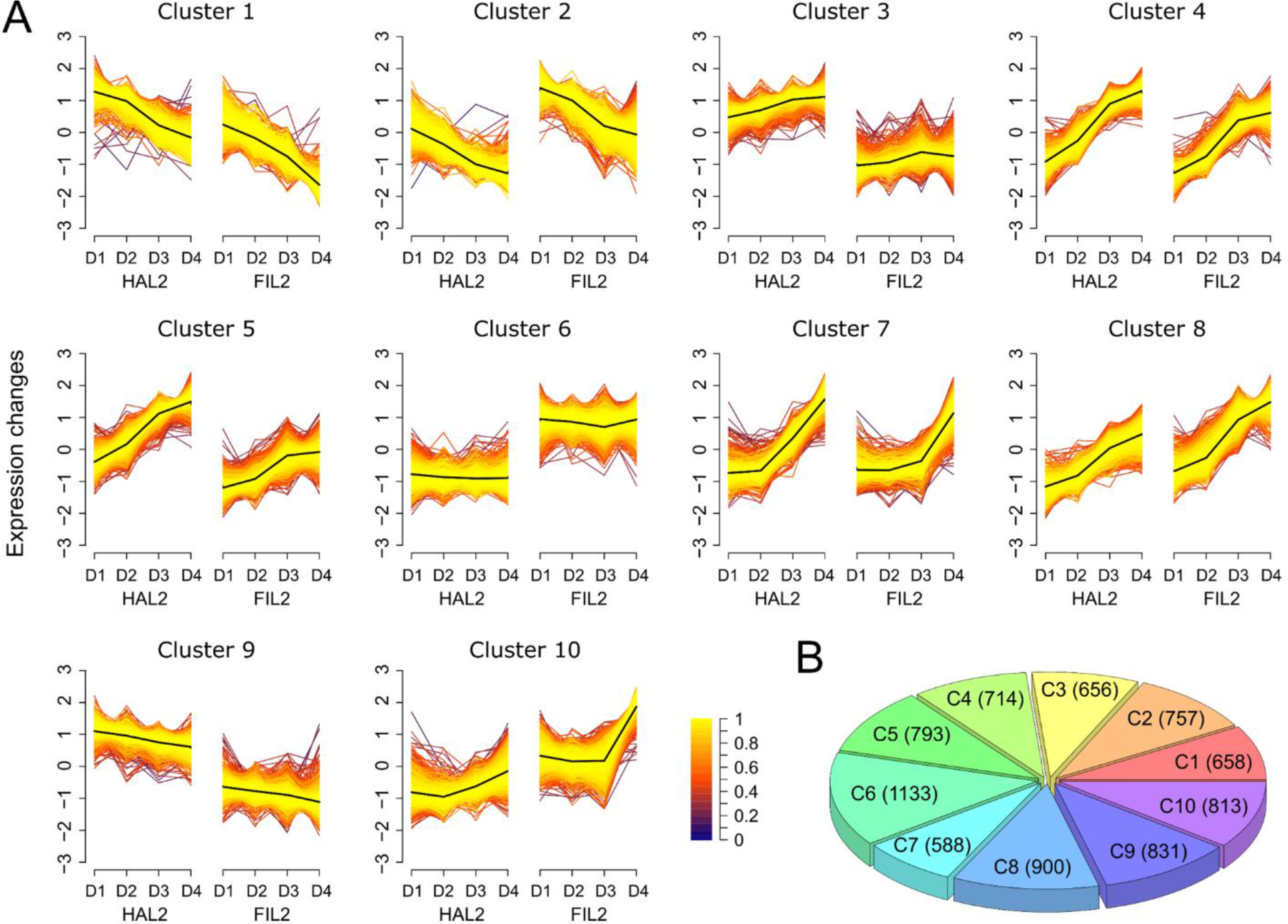
Dynamic patterns of differentially expressed genes during inflorescence development. (A) Fuzzy c-means clustering identified ten distinct dynamic patterns of gene expression divergence. The x axis represents four developmental stages with a side-by-side comparison between HAL2 and FIL2, while the y axis represents variance stabilizing transformed, normalized intensity ratios in each stage. Each line describes the expression pattern of one gene, with the gene’s membership to the cluster represented by the color of the line. The core values for each cluster are plotted in black. (B) The pie chart shows the number of genes in each cluster in (A).

### Global methylome profiles of different *P. hallii* inflorescences

To explore the possible function of DNA methylation in *P. hallii* inflorescence divergence, we generated single-base resolution maps of DNA methylation using bisulfite sequencing. We utilized the same paired inflorescence tissues from HAL2 and FIL2 at the early (D1) and late (D4) stages from our RNA-seq studies for DNA methylation analysis (2 genotypes x 2 developmental stages x 3 biological replicates = 12 libraries). After removal of adapter contaminates and low-quality reads, a total of ∼1.9 billion paired-end reads were generated from two genotypes across two developmental stages each with three biological replicates. We observed a strong mapping bias by performing alignments of the same sequencing reads from all inflorescence samples to both the HAL2 and FIL2 reference genomes. Mapping efficiencies dramatically dropped from ∼75% when aligned to “self” genomes to ∼30% when aligned to incorrect genomes (Supplemental Figure S3, Supplemental Table S3), demonstrating substantial sequence divergence between HAL2 and FIL2. Therefore, we mapped reads from each genotype to their respective genomes for further analysis. We found that approximately 73% of CG sites, 68% of CHG sites, and 55% of CHH sites were covered by at least five uniquely mapped reads across different genotypes and tissues (Supplemental Figure S4). We observed high bisulfite conversion rates, with an average level of 97.5% using a chloroplast control (Supplemental Table S3), that there was little strand differentiation, and the three biological replicates of each sample were highly correlated with each other (*r* > 0.95). These results suggested that our data were reproducible and sufficient for further analysis.

Genome-wide DNA methylation levels analyses revealed that a large proportion of CG (∼66%) and CHG (∼49%) sites have methylated cytosines, while the level of CHH methylation (∼3.1%) was comparatively low (Figure 5D). The genome-wide degree of CG methylation was stable across genotypes and developmental stages, however, we observed a significant difference in non-CG methylation levels (especially in CHH methylation) between genotypes or development stages (*p* < 0.05) (Figure 5D). Global methylation levels revealed that around 32% of CG and 40% of CHG sites had low methylation levels (< 0.2), and about 64% of CG and 32% of CHG sites showed high methylation levels (> 0.8), while CHH site levels were overall very low, with about 97% in the low methylation level category (< 0.2) and less than 0.2% had high methylation levels (> 0.8) (Supplemental Figure S5). The distributions of methylation levels were further compared in three contexts across chromosomes (Figure 5A and 5B; Supplemental Figure S6). We observed a broad hyper CG and CHG methylation region for each chromosome, which is highly negatively correlated with gene density (*r* = -0.974 to -0.976 for CG in all samples; *r* = -0.975 to -0.976 for CHG in all samples) (Figure 5A and 5C; Supplemental Figure S6). These regions are clearly associated with pericentromeres, which have been identified in recent *P. hallii* genomic studies (Lovell et al., 2018). We found a strong positive correlation between CHH methylation levels and gene density in most samples (*r* = 0.573 in HAL2-D1, *r* = 0.741 in FIL2-D1, *r* = 0.626 in FIL2- D4). Intriguingly, this positive correlation was relatively weak in HAL2 inflorescence at the late stage (*r* = 0.199 in HAL2-P4) (Figure 5B and 5C; Supplemental Figure S6).

**Figure 5.**
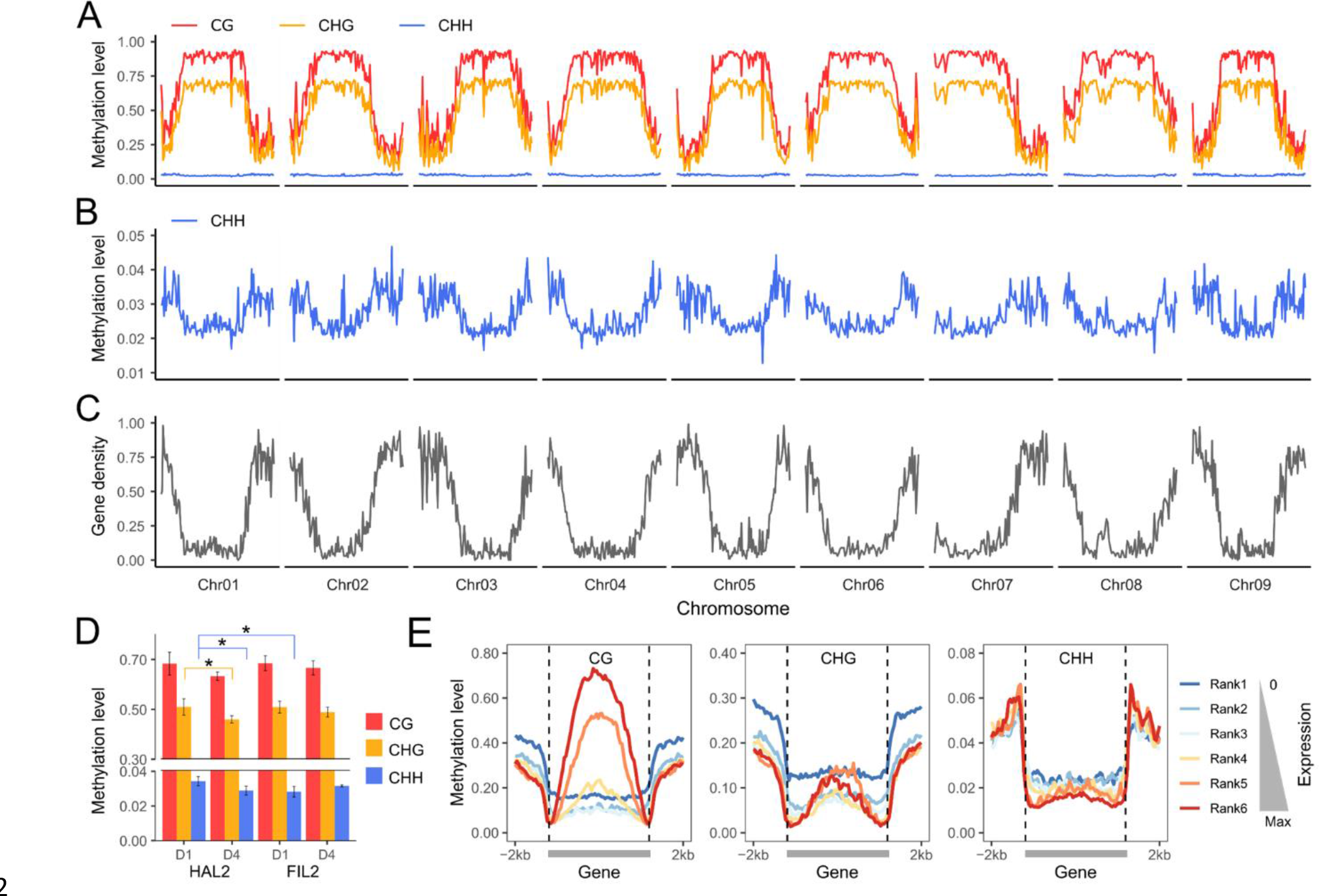
Global DNA methylation profiling and influence of DNA methylation on gene expression during inflorescence development. (A) The distribution of CG, CHG, and CHH methylation levels (mean values of three biological replicates) across the HAL2 chromosomes at D1 inflorescence. (B) The distribution of CHH methylation levels (mean values of three biological replicates) across the HAL2 chromosomes at D1 inflorescence. (C) The distribution of gene density across the HAL2 chromosomes. (D) DNA Methylation levels in different inflorescence tissues and genotypes. (E) Methylation level within gene body and 2 kb flanking regions in CG, CHG, and CHH contexts for the gene sets that are expressed at different levels in HAL2 D1 inflorescence. The average of three replicates was displayed for CG, CHG, and CHH contexts. The data for other genotypes and stages of inflorescences are given in Supplemental Figure S6 and S8.

To understand the relationship between DNA methylation and gene expression, we profiled DNA methylation levels across gene bodies for genes with different expression levels. Genes were divided into six groups based on expression, from a silent rank1 (count = 0) to the highest rank6 (Supplemental Figure S7). We observed that low expression genes, especially the genes with no expression in rank1, have higher CG and CHG methylation level at the promoter and the 3’ end regulatory regions (Figure 5E and Supplemental Figure S8). Notably, we found that high expression genes, especially the genes in rank6 and rank5, have higher CG methylation level at the gene body regions (Figure 5E and Supplemental Figure S8). These patterns were observed across both genotypes and all development stages (Supplemental Figure S8), suggesting the diverse roles of DNA methylation in *P. hallii* development. Methylation in promoter regions was associated with transcriptional silencing while methylation in gene-body regions was positively correlated with gene expression levels.

### Differential DNA methylation between different *P. hallii* inflorescences

To determine differentially methylated regions, we compared methylation levels across different genomic regions in one-to-one orthologous genes between two genotypes (HAL2 vs FIL2) or two development stages (D1 vs D4). A gene with a significantly different proportion of methylation in any of the three methylation contexts across at least one annotated feature was considered a differentially methylated gene (DMGs) (*q* < 0.01, methylation level change > 0.1, see method for details). Overall, we detected 12,053 (51.3%) DMGs between HAL2 and FIL2 across inflorescence development. 10,141 (43.1%) genes are differentially methylated at the earliest stage and 9,134 (38.8%) genes are differentially methylated at the late stage (Figure 6, Supplemental Figure S9, Supplemental Table S4, and Supplemental Table S5). In total, we discovered 7755 (33%) DMGs in the CG context, 5784 (24.6%) DMGs in the CHG context, and 2783 (11.8%) DMGs in the CHH context at the early stage (Figure 6). Similarly, a total of 6744 (28.7%) DMGs in the CG context, 5228 (22.2%) in the CHG context, and 2555 (10.9%) DMGs in the CHH context were detected at the late stage (Supplemental Figure S9). These results implicate DNA methylation in inflorescence divergence in *P. hallii*. Notably, our analysis revealed that most of DMGs were differentially methylated at the promoter, with a higher number resulting from FIL2 hypermethylation in all three methylation contexts (Figure 6 and Supplemental Figure S9). Moreover, we detected a large number of DMRs occurred in the untranslated region, especially the 3-prime end (Figure 6 and Supplemental Figure S9). In addition, a smaller number of DMRs were found in CDS or intron (Figure 6 and Supplemental Figure S9). These results suggest that the flanking regions (promoter and 3UTR regions) are more frequently differentially methylated than the regions within genes (exons and introns). Further, we compared the methylation levels between early and late stages of inflorescences in each genotype. We only detected 913 (3.9%) DMGs associated with HAL2 inflorescence development (Supplemental Figure S10 and Supplemental Table S6) and 740 (3.1%) DMGs associated with FIL2 inflorescence development (Supplemental Figure S11 and Supplemental Table S7), suggesting that methylation levels are relatively stable during inflorescence development in *P. hallii*.

**Figure 6.**
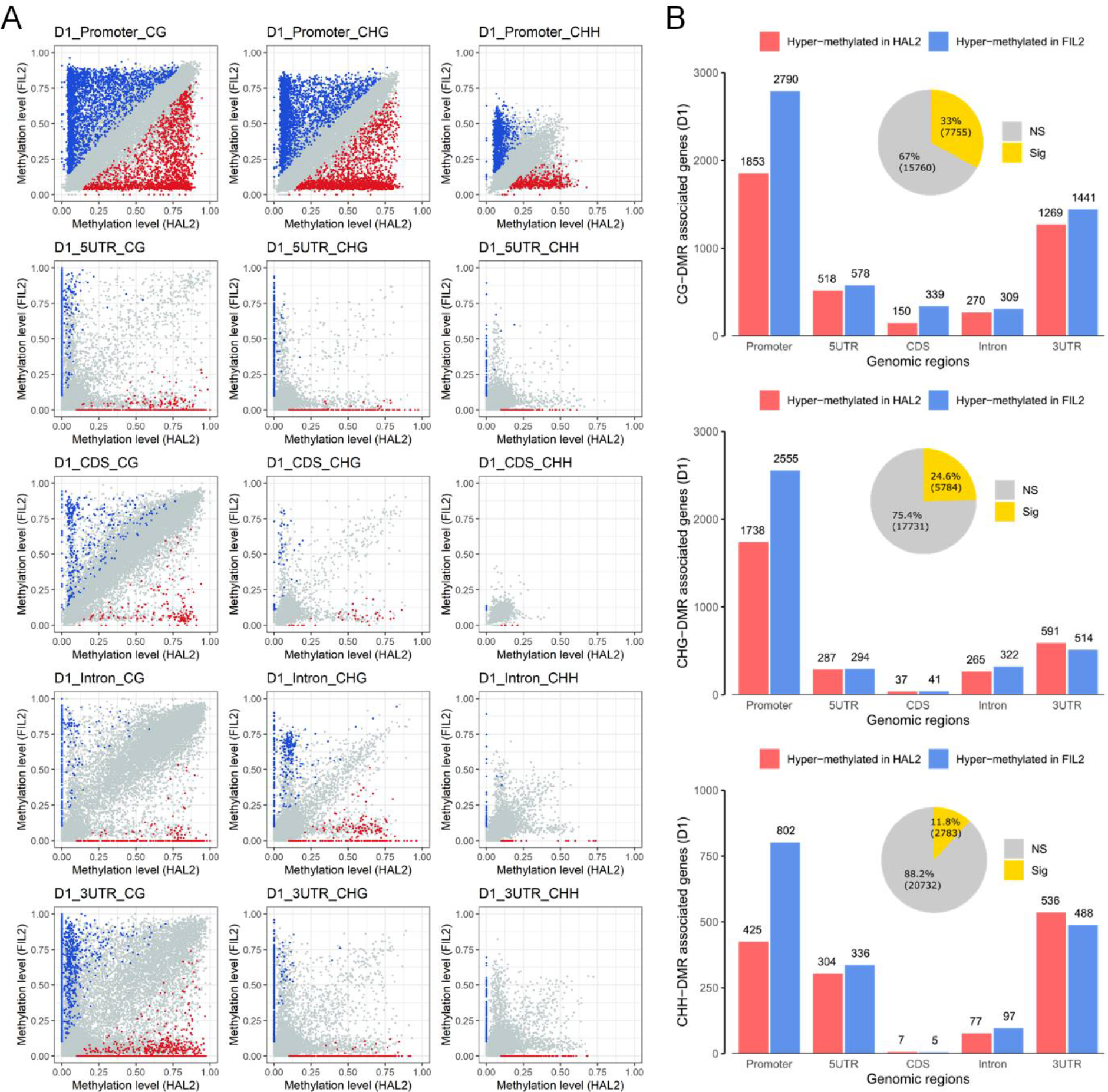
Differential DNA methylation regions between HAL2 and FIL2 inflorescences. (A) Pairwise comparisons of methylation levels from one-to-one orthologous gene pairs between HAL2 and FIL2 D1 inflorescence in CG, CHG, and CHH contexts across five different genomic features. Blue dots represent genes with significant hypermethylation in FIL2, while red dots represent genes with significant hypermethylation in HAL2. Grey dots represent genes with no significant methylation difference. (B) Number of differentially methylated genes between HAL2 and FIL2 D1 inflorescence in CG, CHG, and CHH contexts across five different genomic features are shown in bar plots. The total number of differentially methylated genes in each context is shown in pie chart. The data for the comparison between HAL2 and FIL2 at D4 inflorescence and the comparison between D1 and D4 inflorescence in both HAL2 or FIL2 background are given in Supplemental Figure S9, S10 and S11.

### The patterns of DMR-associated DEGs evolution during inflorescence divergence

To understand the role of DNA methylation in driving the expression of genes involved in the inflorescence diversity, we identified DMR-associated DEGs between the ecotypes by joining the results of our methylome and transcriptome datasets. A total of 2,057 and 2,432 genes at D1 and D4 stages, respectively, were identified as DMR-associated DEGs between HAL2 and FIL2 inflorescences (Supplemental Figure S12), suggesting that almost half of DEGs were associated with the methylation changes (49.7% in D1 and 43.0% in D4). Among them, 1,672 genes in CG context, 1,193 genes in CHG context, and 607 genes in CHH context were detected as DMR-associated DEGs at D1 stage (Supplemental Figure S12). Similarly, 1,844 genes in CG context, 1,392 genes in CHG context, and 726 genes in CHH context were detected as DMR-associated DEGs at D4 stage (Supplemental Figure S12). Overall, DMR-associated DEGs with CG context differential methylation were more abundant than other sequence contexts. We observed that a larger fraction of DMR-associated DEGs had differences in methylation at the flanking regulatory regions, especially the promoter and 3UTR regions (Figure 7 and Supplemental Figure S13), suggesting the importance of these regions. Intriguingly, we did not observe a simple pattern between the direction of differential methylation and differential gene expression at both stages of inflorescence development (Figure 7 and Supplemental Figure S13). The trend between direction of differential methylation and differential gene expression could be positive or negative in all three sequence contexts located in the five different genic regions (Figure 7 and Supplemental Figure S13). Similar findings are also reported in previous studies (Rajkumar et al., 2020), suggesting the complex role DNA methylation plays in gene expression regulation.

**Figure 7.**
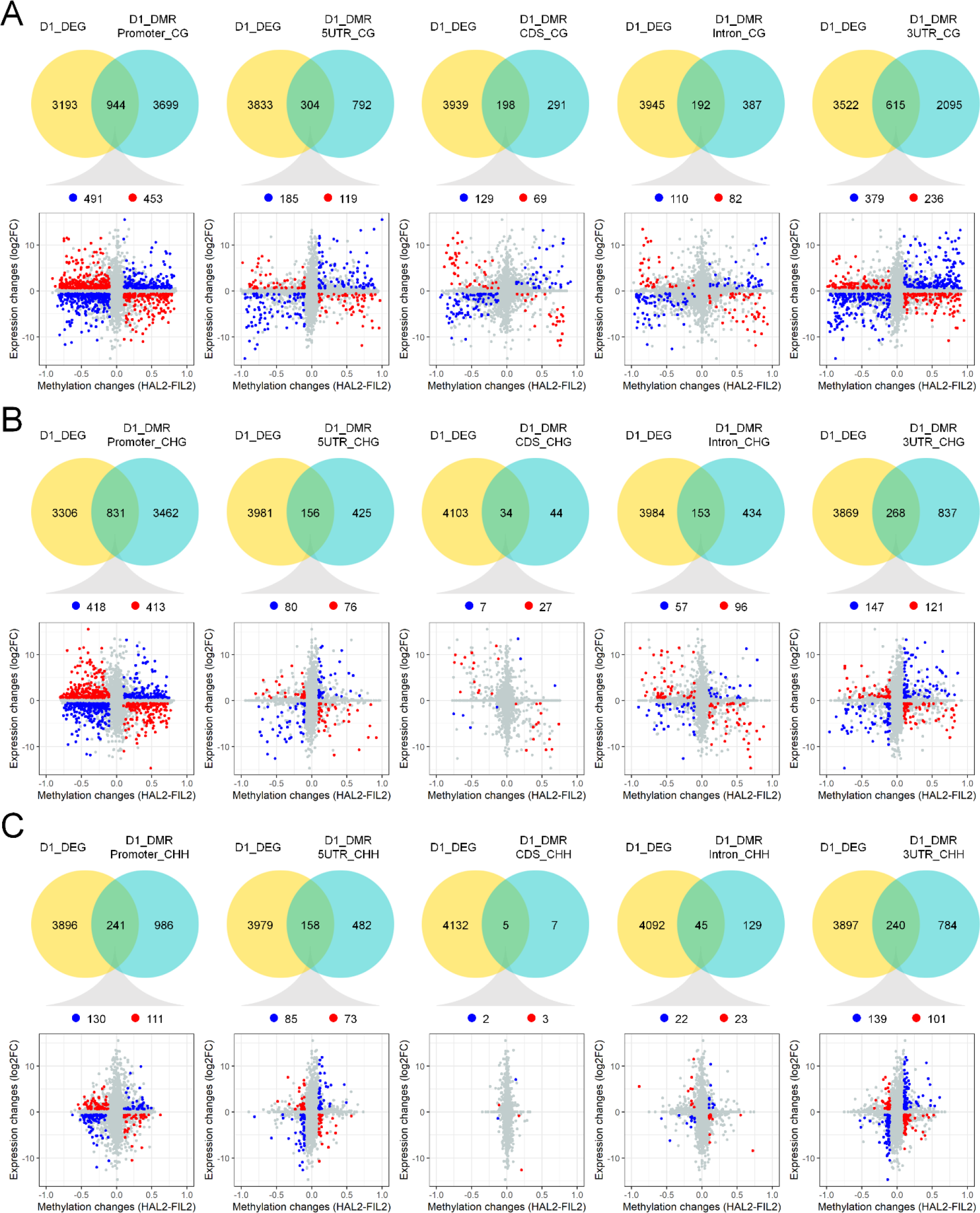
Association of differentially methylated genes with differentially expressed genes. Venn diagrams depicting the number of differentially expressed genes (yellow circle, DEG) and DMR-associated genes (blue circle, DMR) between HAL2 and FIL2 D1 inflorescence in CG (A), CHG (B), and CHH (C) contexts across five different genomic features. Two-dimensional scatter plots depict the association of differentially expressed genes and differentially methylated regions in CG (A), CHG (B), and CHH (C) contexts across five different genomic features. The x axis represents relative gene expression change (log2fold change), while the y axis represents relative methylation change (HAL2 subtract FIL2). The data for the association of differentially methylated genes with differentially expressed genes at D4 inflorescence are given in Supplemental Figure S13.

To explore the pattern of protein evolution associated with DMR-associated DEGs, we compared the *Ka*/*Ks* ratio (non-synonymous substitutions per non-synonymous sites/synonymous substitutions per synonymous sites) for HAL2 and FIL2 gene pairs between DMR-associated DEGs and the genome-wide pattern for one-to-one orthologous genes. The *Ka*/*Ks* ratio of DMR-associated DEGs and one-to-one orthologous genes centered around a peak at 0.55 and 0.46 (Figure 8A), respectively. No statistically significant difference of the *Ka*/*Ks* ratio distribution was observed between DMR-associated DEGs and the genome-wide backgrounds. We observed that only 348 (∼12.8%) of DMR-associated DEGs pairs have *Ka*/*Ks* ratios > 1 (Figure 8B), suggesting that the majority of DMR-associated DEGs are evolving under purifying selection.

**Figure 8.**
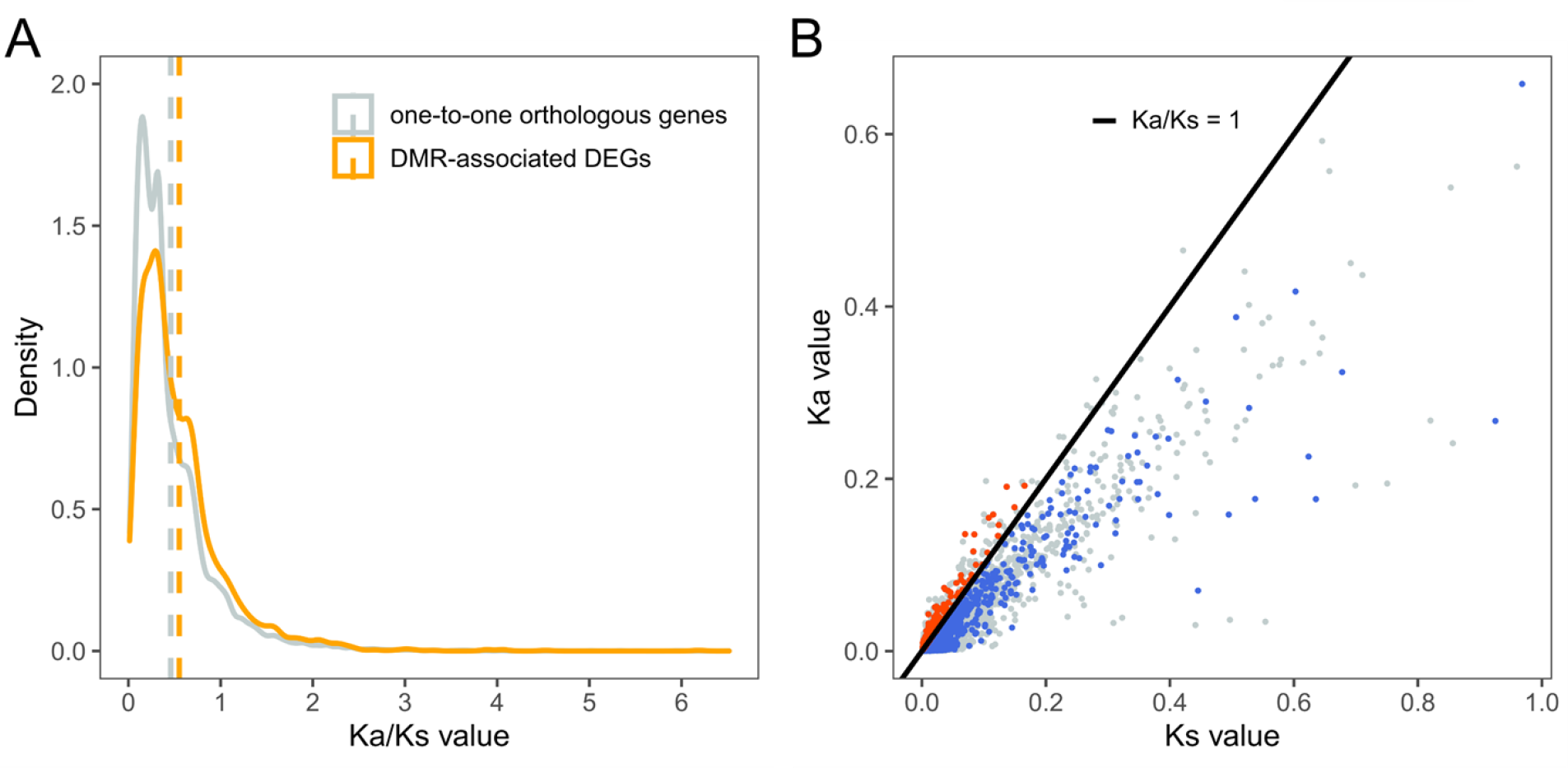
K_a_/K_s_ distribution of DMR-associated DEGs gene pairs. (A) and (B) The Ka/Ks values distribution of gene pairs from DMR-associated DEGs and one-to- one orthologous genes between HAL2 and FIL2. (A) The mean values are indicated by the dashed line. (B) The solid black line marks Ka/Ks = 1. The red dots mark the DMR-associated DEGs with Ka/Ks ratio larger than 1, while the blue dots mark the DMR-associated DEGs with Ka/Ks ratio less than 1. The grey dots represent all one-to-one orthologous genes.

Further, we analyzed the differential methylation and expression of candidate genes involved in inflorescence and seed development processes, based on reports from other grass model systems. We identified candidates from DMR-associated DEGs including, for example, the homologs of *SPL14* (PhHAL.6G248800) (Miura et al., 2010), *DEP1* (PhHAL.2G271000) (Huang et al., 2009), *Ghd7.1/PRR37* (PhHAL.2G086000) (Yan et al., 2013), and *MADS1* (PhHAL.9G564100) (Khanday et al., 2013) (Figure 9A and 9B). These candidates are hub genes sharing the same regulation network among key transcription factor families (*SPL* and *MADS*), G-protein signaling, and flowering time pathways (Lu et al., 2013; Liu et al., 2018; Liu et al., 2018), which determine both inflorescence and seed development in many grass systems. We observed FIL2 hypermethylation patterns, especially for the CG context within 3UTR region, for the homolog of *SPL14*, and HAL2 hypermethylation patterns in CG and CHG contexts within promoter regions for the homologs of *DEP1*, *Ghd7.1/OsPRR37*, and *OsMADS1* (Figure 9A). The homologs of *DEP1* and *SPL14* had a similar expression pattern, with HAL2 showing consistently greater gene expression relative to FIL2 and with both genotypes showing decreased expression from D1 to D4 stages (Figure 9B). The homolog of *MADS1* had greater expression in FIL2 relative to HAL2 across different stages, while the *Ghd7.1/OsPRR37* homolog had a heterochronic change with greater gene expression in FIL2 relative to HAL2 at the early stages (Figure 9B). Differential methylation and expression of these candidate genes may play an important role in the evolution of *P. hallii* inflorescences.

**Figure 9.**
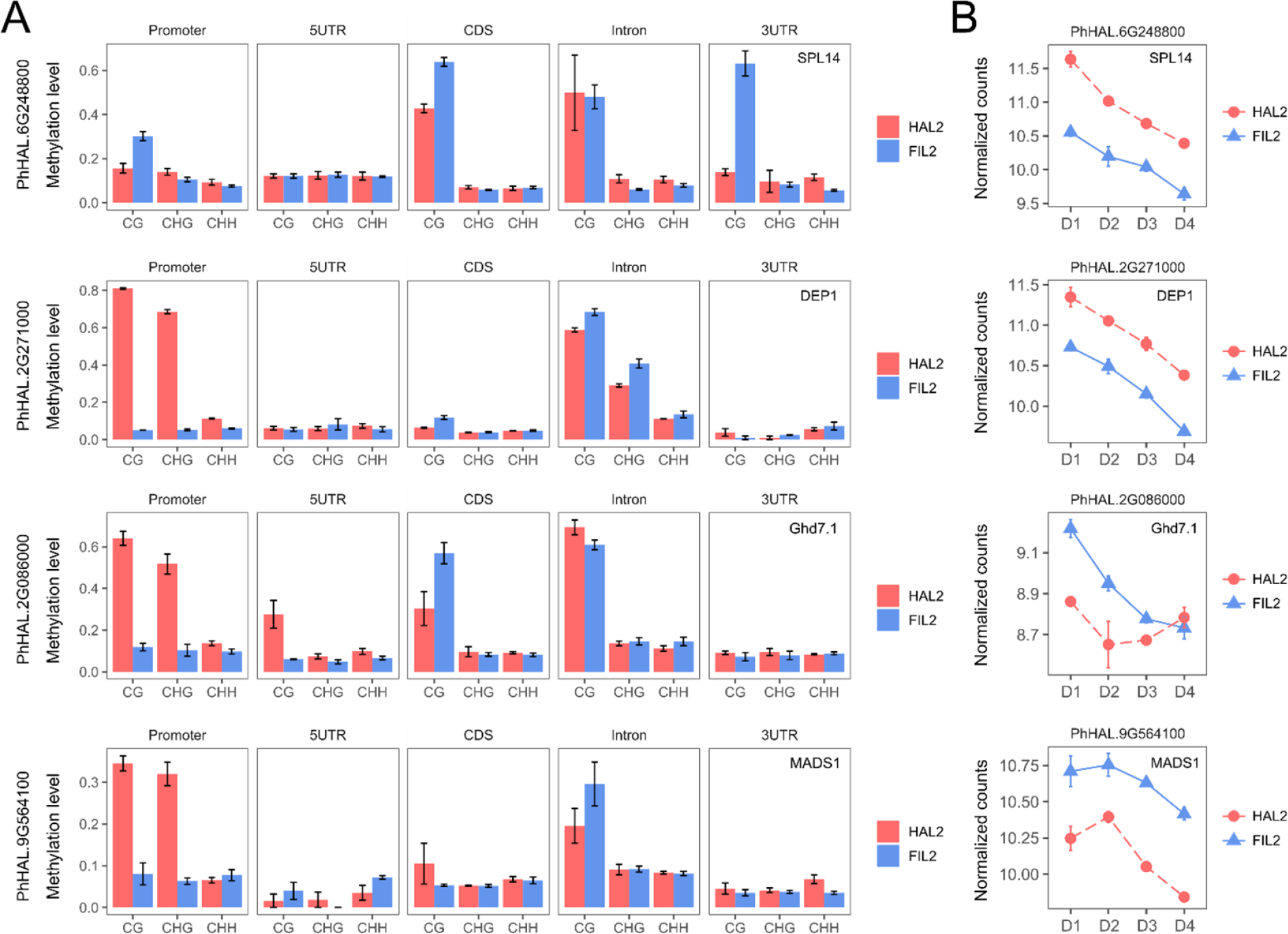
Differential methylation and differential expression patterns of candidate genes involved in the inflorescence and seed development. (A) Differential methylation patterns of CG, CHG, and CHH contexts across five different genomic features. In all panels, the bar plots and error bars are the average values and SE, respectively, based on methylation level from three replicates. (B) Differential expression patterns of differentially methylated genes in (A). The x axis represents four developmental stages, while the y axis represents normalized counts using variance stabilizing transformation in DEseq2. In all panels, the points and error bars are the average values and SE, respectively, based on normalized counts of three RNA-seq replicates.

## Discussion

For members of the Poaceae, the branching system of the inflorescence determines the number of flowers formed and the size and number of seeds, which are fundamental life-history traits in wild species and critical economic traits in crop yield potential. Genome-wide gene expression and DNA methylation analyses are now widely used to study the genetic and epigenetic mechanisms of inflorescence development from a variety of important crops (Furutani et al., 2006; Wang et al., 2010; Li et al., 2012; Eveland et al., 2014; Feng et al., 2017; Parvathaneni et al., 2020; Sun et al., 2020). Despite ever-increasing knowledge of grass inflorescence development, an effective model of inflorescence patterning from wild species without a domestication history is still lacking. *P. hallii* is a native perennial C4 grass with a highly diverse and complex inflorescence with striking divergence between ecotypes adapted to contrasting habitats (Lowry et al., 2015; Palacio-Mejia et al., 2021). To date, *P. hallii* has been developed as a complementary diploid model in parallel with domesticated crops and other C4 perennial grasses (Lovell et al., 2018). A systematic comparison of inflorescence transcriptome and DNA methylome for *P. hallii* ecotypes should provide novel insights into the molecular mechanisms leading to their divergent inflorescences.

### Expression divergence of *P. hallii* inflorescence development

Previous transcriptomic studies have revealed the regulatory modules of young inflorescence development in major crops including rice, maize, and wheat (Furutani et al., 2006; Wang et al., 2010; Eveland et al., 2014; Harrop et al., 2016; Feng et al., 2017). These studies provide resources to identify potential targets for genetic engineering and overall crop improvement. However, most of these studies performed experiments within a single genetic background, and therefore provide limited information about the evolution of gene regulatory networks associated with traits under selection. Using *P. hallii* as a model wild perennial grass, we designed a two-factor factorial experiment to understand gene expression divergence and development. Despite only ∼1.08 Mya of divergence (Lovell et al., 2018), we demonstrated that the vast majority of genes exhibited significant expression divergence between HAL2 and FIL2 inflorescence or significant expression level changes across development. We detected a large number of genes with significant interactions between development and genotype, suggesting the importance of heterochronic changes in gene expression during the development of inflorescence evolution in *P. hallii*. Heterochronic change is an alteration in the timing of developmental programs during evolution, which are known to contribute to the evolution of inflorescence architecture (Buendia-Monreal and Gillmor, 2018). In grasses, the inflorescence branching systems are determined by the timing of phase transition from the branch meristem to the spikelet meristem (Kyozuka et al., 2014). Delays in the spikelet meristem specification result in more complex and larger inflorescences (Kyozuka et al., 2014). In our study, we observed heterochronic shifts for the homologs of *Ghd7.1*/*OsPRR37* and a number of genes involved in auxin, cytokinin, and gibberellin pathways. *Ghd7.1*/*OsPRR37* is a core circadian gene controlling flowering time, inflorescence architecture, and adaptation (Yan et al., 2013). Hormonal regulation of meristem initiation, size, and determinacy play a key role in inflorescence development (Barazesh and McSteen, 2008). Therefore, the heterochronic changes in expression of these genes across development could shift aspects of meristem differential or patterns of cell division, which result in the evolution of the reduced structure of HAL2 inflorescence relative to the open architecture of FIL2 inflorescence.

Previous studies have highlighted the importance of transcription factors at the early stage of inflorescence development and their conserved roles between rice and other crops (Harrop et al., 2016; Zhu et al., 2018). For example, *SPL14* and additional *SPL* genes, such as *SPL7* and *SPL17*, were detected at the early stages of inflorescence development in different crops (Wang et al., 2010; Feng et al., 2017). The functional validation of *SPL* genes from rice and maize revealed an evolutionarily conserved relationship between the expression level of *SPL* genes and the branching structure of the inflorescence (Du et al., 2017). In our study, the *P. hallii SPL7*, *SPL14*, and *SPL17* homologs showed higher expression in HAL2 relative to FIL2 and existed in cluster 1 with the tendency to decrease across developmental gradients. Similarly, *MADS-box* genes control inflorescence branching systems via the regulation of meristem identification and development in other grasses (Liu et al., 2013; Kyozuka et al., 2014). We found that a large number of *MADS* homologs, including *MADS1*, *MADS14*, and *MADS18*, were more highly expressed in the FIL2 relative to HAL2 with a general declining pattern of expression with development in cluster 2. These expression patterns suggest an important role of *SPL* and *MADS* genes in the divergence of *P. hallii* inflorescence architecture determination.

### DNA methylation and the evolution of inflorescence development in *P. hallii*

In this study, we performed whole-genome bisulfite sequencing to understand the role of DNA methylation during inflorescence development and architecture divergence. As the first inflorescence methylome study in *P. hallii*, we found that the overall methylation levels from tissues across different genotypes and development stages were similar in each context and the proportion of methylated cytosines in CG, CHG, and CHH contexts are 66%, 49%, and 3.1%. For comparison, values reported in *Arabidopsis* are 30.5% for CG, 10.0% for CHG, and 3.9% for CHH, in rice are 58.4% of CG, 31.0% of CHG, and 5.1% for CHH, and in maize are 86% of CG, 74% of CHG, and 5.4% for CHH (Niederhuth et al., 2016). These results suggested that *P. hallii* has an intermediate level of DNA methylation. Coincident with previous findings, we observed a strong positive correlation between CG and CHG methylation with gene density, suggesting the role of DNA methylation in the establishment and maintenance of centromeric and pericentromeric heterochromatin regions. On the other hand, we found a positive correlation between CHH methylation and gene density. Previous studies have shown that global hypermethylation in CHH context in rice reproductive organs might function as the protective mechanism for the genome. Our analysis of the relationship between gene expression and DNA methylation level suggested that non-expressed and low-expressed genes showed higher CG and CHG methylation levels in their proximal regulatory regions, while genes expressed at high levels were highly CG methylated within their gene body regions. These patterns are similar to recent results reported in chickpea and papaya (Rajkumar et al., 2020; Zhou et al., 2020), suggesting the conserved antagonistic role of CG methylation in gene expression regulation at the regulatory and gene body regions.

Previous studies have shown that mapping bias to a single genome can introduce clear and substantial quantification bias in the identification of DMR (Wulfridge et al., 2019). Unfortunately, most methylome studies align to a single reference genome to identify DMR between different genotypes due to the limitation of genomic resources (Li et al., 2012; Rajkumar et al., 2020). Here, we observed a dramatic drop in mapping efficiencies from alignments to individual genomes (HAL2 to HAL2 and FIL2 to FIL2) (∼75%) compared to alignments to divergent genomes (HAL2 to FIL2 and FIL2 ->HAL2 (∼30%). After mapping reads to their own individual genome references, we found that more than half of one-to-one orthologous genes (12,053) are differentially methylated in at least one feature of genomic regions between HAL2 and FIL2 across inflorescence development. This degree of widespread natural variation in DNA methylation was also observed in a diverse panel of *Arabidopsis* and maize (Kawakatsu et al., 2016; Xu et al., 2020). Interestingly, previous studies showed that differential methylation primarily occurs within gene body regions (Rajkumar et al., 2020). However, we observed that flanking regulatory regions, including promoter and 3UTR, are more frequently differentially methylated than the regions within gene body regions. One explanation for this conflicting pattern could be difference in alignment strategy, as most of these studies mapped reads from different genotypes to one genome reference and probably induced quantification bias in the highly variable regulatory regions. Although tissue or developmental stage specific methylation patterns have been mentioned in some studies (Huang et al., 2019), we only detected a few DMGs between two inflorescence tissues. Considering that a large number of one-to-one orthologous genes are differentially expressed with significant development effects, these genes are most likely driven by other factors but not by the methylation difference across inflorescence development. Finally, we detected more than 3,000 DMR-associated DEGs between HAL2 and FIL2 across inflorescence development. The relationship between the direction of differential methylation in different sequence contexts and differential gene expression is not simple, including both positively and negatively correlated patterns. This complex pattern was also observed from a recent study of seed development in chickpea (Rajkumar et al., 2020). Although recent evidence from population level studies suggested the selection on DNA methylation could be weak, differential methylation of key development genes (e.g. *vgt1* and *tb1* in maize) are associated with phenotypic variation (Xu et al., 2020). Therefore, function validation of DMR-associated DEGs identified in the present study will help to understand the inflorescence divergence in *P. hallii*.

### Molecular regulation of seed size-number trade-off in *P. hallii*

To maximize fitness, plants are assumed to allocate their resources optimally between producing fewer large seeds versus more small seeds, termed the seed size-number trade-off (Venable, 1992). Although this fundamental life-history trade-off is widely accepted from an ecological and evolutionary perspective, strong evidence of its genetic and molecular basis is scarce and inconclusive (Dani and Kodandaramaiah, 2017). At the phenotypic level, a strong negative correlation between seed size and seed number has been frequently observed within and across plants species (Sadras, 2007). However, at the genetic level, QTL analysis of seed size and seed number in *Arabidopsis*, suggests that the genetic factors controlling these two traits are mostly nonoverlapped (Gnan et al., 2014), suggesting a minor role of seed size-number trade-off in its evolutionary history. Here, we observed that HAL2 produced many fewer large seeds compared to FIL2. We hypothesize that this variation in seed number is driven by the difference in inflorescence branching structure, and a potential seed size-number trade-off in *P. hallii* could result from long-term adaptation to their local environmental stress. Further studies of natural variation of traits related to seed size-number trade-off in *P. hallii* along with genetic mapping of these traits may help to understand the life-history strategies of this wild perennial grass.

Numerous key genetic and molecular switches have been identified to regulate inflorescence patterning by determining the fate of reproductive meristem (Zhang and Yuan, 2014; Li et al., 2021). Major regulators and signaling pathways have been revealed to determine the seed size by the integration of cell proliferation and expansion in the embryo, endosperm, and seed coat (Li et al., 2019). These advances provide an opportunity to understand the interaction and coordination of inflorescence and seed development. It has been noted that G-protein signaling is one of the critical mechanism involved in both inflorescence architecture and seed size regulation (Li et al., 2019; Li et al., 2021). Recent studies have identified multiple subunits of the heterotrimeric G proteins that regulate seed size and shape across different species (Li et al., 2012; Sun et al., 2018). Among them, *DEP1* is a heterotrimeric Gγ subunit (Sun et al., 2018), which was initially isolated as a major yield QTL controlling inflorescence architecture in rice (Huang et al., 2009). A dominant allele of *DEP1* enhanced meristematic activity and cell proliferation, resulting in the increased branch number and grain number but decreased grain weight (Huang et al., 2009). In this study, we identified the homolog of *DEP1* as a DMR-associated DEG, suggesting the potential role of *DEP1* in the regulation of seed size-number trade-off in *P. hallii*. Intriguingly, several *DEP1* related genes, including *SPL14*, *MADS1*, and *Ghd7.1/PRR37*, were also differentially methylated and expressed between HAL2 and FIL2 inflorescence. Chromatin immunoprecipitation- sequencing analysis revealed that *SPL14* could directly bind to the promoter of *DEP1* for the regulation of inflorescence architecture (Lu et al., 2013). Protein analysis revealed that *DEP1* determines seed size through interaction with *MADS1* (Liu et al., 2018). *Ghd7.1/OsPRR37* is an upstream regulator that regulates the expression of *MADS1* (Liu et al., 2018). Evolutionary analyses of DMR-associated DEG candidate genes suggest that their protein functions are largely conserved, with the vast majority exhibiting signatures of purifying selection. This may indicate that adaptive evolution and divergence in the *P. hallii* inflorescences has largely occurred through regulatory evolution in combination with epigenetic modifications. Although the epigenetic mechanisms responsible for gene regulation are still unclear, the observed divergence in DNA methylation and gene expression of these candidate genes might play a crucial role in the seed size-number trade-off and inflorescence divergence between the upland and lowland ecotypes in *P. hallii*.

## Materials and Methods

### Plant materials and sample collection

*P. hallii* genotypes, HAL2 (*P. hallii var. hallii*, upland ecotype) and FIL2 (*P. hallii var. filipes*, lowland ecotype), were grown in a growth chamber at University of Texas at Austin with 26 °C day/ 22 °C night temperature and 12-hr photoperiod. Plants were grown in 3.5 in. square pots with a 6:1:1 mixture of Promix:Turface:Profile soil. The first fully emerged inflorescence was photographed and used to measure the primary branch number, secondary branch number, and spikelet number as previously described (Wang et al., 2015). The seeds were harvested after maturity and dried at a temperature of 37 °C until the seed weight was stable. The dried seeds were photographed and weighed for the 100-seed weight (mg) value. Phenotypic values are averages from eight replicates showing uniform growth. Young panicle tissues were collected under a dissection microscope and the developmental stages were determined according to the lengths (0.1-0.2 cm for D1 stage, 0.5-1 cm for D2 stage, 4.5-5.5 cm for D3 stage, and 9-11 cm for D4 stage). Tissues for D1 and D2 stage were taken from at least fifty plants and pooled for each biological replicate. Tissues for D3 and D4 stage were taken from at least fifteen plants and pooled for each biological replicate. All samples were harvested at 17:00-18:00 of the day and immediately flash frozen in liquid nitrogen. Three biological replicates for each stage were stored at -80 °C for DNA and RNA extraction.

### Sequence analysis

19,332 one-to-one orthologous genes were identified in a previously published *P. hallii* genomic study (Lovell et al., 2018). The synonymous substitution rates (Ks), non-synonymous rates (Ka) and non-synonymous to synonymous substitution ratios (*Ka*/*Ks*) of all one-to-one orthologous gene pairs of HAL2 and FIL2 were estimated by using the “simple Ka/Ks calculator” function from TBtools (Chen et al., 2020).

### RNA extraction and RNA-seq library preparation

For RNA preparation, inflorescence samples from four development stages were homogenized to fine powder using a pre-chilled mortar and pestle under liquid nitrogen. Total RNA was isolated using the TRIzol kit (Invitrogen) and samples were treated with DNase I (Invitrogen) to remove contaminating genomic DNA. RNA-Seq libraries were prepared and sequenced in the Department of Energy Joint Genome Institute (Walnut Creek, CA). Briefly, the integrity and concentration of the RNA preparations were checked initially using Nano-Drop (Nano-Drop Technologies) and then by BioAnalyzer (Agilent Technologies). Total RNA-Seq libraries were prepared using Illumina’s TruSeq Stranded mRNA HT sample prep kit utilizing poly-A selection of mRNA. Sequencing was performed on the Illumina HiSeq 2500 platform using HiSeq TruSeq SBS sequencing kit, following a 2 × 150 indexed run recipe.

### RNA-Seq data analyses

Paired-end RNA-Seq 150-bp reads were quality trimmed (*Q* ≥ 25) and reads shorter than 50 bp after trimming were discarded. High-quality filtered reads were aligned to their own reference genomes, *Panicum hallii* HAL v2.1 and *Panicum hallii* v3.1 (https://phytozome-next.jgi.doe.gov/), using GSNAP with a maximum of four mismatches. The HTseq-count was used to generate raw gene counts, and only reads that uniquely mapped to one annotated gene were counted. To filter the genes with low expression and compare the diverged transcriptome assemblies, only one-to-one orthologous genes with counts-per-million above 0.5 (correspond to a count between 8-10 for different library sizes) in at least three samples were retained for further analysis. Principal component analysis and specific gene expression patterns were performed with *vst* normalized expression counts and visualized using the ggplot2 package. Differentially expressed genes with main effects and developmental-specific effects were determined as previously described (Weng et al., 2019). Briefly, to study the additive and interaction effects of genotype and development stage, we determined differential gene expression using statistical testing via likelihood ratio tests in DEseq2. We used a factorial linear model to test the following: (a) genotype additive effect by comparing the difference in deviance between the two-factor additive model (Genotype + Development stage) and a reduced model (Development stage) formula; (b) the effect across development stages by comparing the difference in deviance between the two-factor additive model (Genotype + Development stage) and a reduced model (Genotype); and (c) interaction effect by comparing the difference in deviance between the full model (Genotype + Development stage + Genotype × Development stage) and an additive reduced model (Genotype + Development stage). Multiple testing was controlled by *q*-value transformation of likelihood ratio test *p*-value and genes with expression divergence were determined by significant genotype and/or interaction effects (*q* < 0.01). To further study developmental stage-specific effects, we conducted a linear model fit to a set of four HAL2-FIL2 contrasts with genes exhibited genotype and/or interaction effects, one at each development stage, through a custom contrast analysis pipeline in DEseq2 with the calculation of log2-fold change values and adjusted *p*-value (*q*-value). The profile of genotype predominant genes with developmental stage-specific information was plotted with the UpSetR package. The *vst* transformed counts of genes with strong expression variation were used for soft clustering with the Mfuzz package.

### DNA extraction and bisulfite sequencing library preparation

A CTAB-based protocol was used for DNA extraction from D1 and D4 inflorescence samples of both genotypes (HAL2 and FIL2). The quality of DNA was determined by running on a 1.0 % agarose gel electrophoresis and quantified via Nano-Drop (Nano-Drop Technologies). DNA methylome libraries were prepared from 1ug of genomic DNA and underwent bisulfite treatment using NextFlex Bisulfite-Seq Kit. The resulting bisulfite-converted DNA was PCR- amplified and ligated to adapters, with barcodes. Amplified fragments were purified using the 1.8 x AMPure XP a bead cleaning to remove the small fragments. The libraries were checked for size and concentration using the Agilent Bioanalyzer instrument, followed by sequencing on the Illumina HiSeq 2500 platform at HudsonAlpha Institute.

### Bisulfite sequencing data analyses

Paired-end bisulfite sequencing 150-bp reads were trimmed using Trim Galore with default options to remove low-quality reads and adaptor sequences. To avoid mapping bias induced by divergent reference genomes, high-quality filtered reads were aligned to their own respective reference genomes, *Panicum hallii* HAL v2.1 and *Panicum hallii* v3.1 (https://phytozome-next.jgi.doe.gov/), using Bismark with options --bowtie2 –bam (Krueger and Andrews, 2011).

Reference genomes index files for HAL2 and FIL2 were generated from their corresponding FASTA files using the bismark_genome_preparation function. After removing duplications with the deduplicate_bismark function, BAM output files were sorted in preparation for methylation extraction using Samtools. Genome-wide cytosine reports were obtained using the bismark_methylation_extractor with options -p -ignore 5 -ignore r2 5 -ignore 3prime 2 -ignore 3prime r2 -no_overlap –comprehensive -CX. This report was used to generate the read coverage, global methylation level, and distribution of methylation level using ViewBS (Huang et al., 2018). Reads mapped to unmethylated chloroplast genome were used to calculate the frequency of cytosine conversion. For DMR analysis, only cytosines that were covered by at least five reads were kept for downstream analysis. We studied differential methylation between ectotypes or developmental stages using a genomic feature approach. We defined genomic regions to include promoter regions from 2-kb upstream of the transcription start site, 5’-untranslated region (5UTR), protein-coding region (CDS), intron, and 3’-untranslated region (3UTR) based on the annotation of gene structure from the existing *P. hallii* genome GFF files for each respective genome. The methylation level in each genomic region was measured as the average of the proportion of all methylated cytosines in that region. The methylation levels of different genomic regions from one-to-one orthologous genes were extracted and used for DMR analysis. For each DMR contrast, we performed a student *t*-test and calculated the *q*-value using qvalue package to control for the large number of statistical tests. We calculated the methylation changes by subtracting average methylation proportions from HAL2 to FIL2. A cut-off of < 0.05 *q*-value and > 0.1 methylation change were used to identify significant DMR across five genomic regions and three methylation contexts. In a small number of cases, methylated cytosines were detected for one level of a contrast but not for the other (e.g., methylation was observed in only one ecotype or developmental state for a feature). For these genes/regions we simply used a cut-off > 0.1 methylation proportion change to identify a significant DMR.

## Accession numbers

The RNA sequencing and bisulfite sequencing data generated in this study have been deposited with NCBI at XXX under series accession numbers XXX and XXX, respectively.

## Supplemental data

Supplemental Figure S1. Determination of the number of cluster cores.

Supplemental Figure S2. Expression pattern of candidate genes in different clusters.

Supplemental Figure S3. Mapping efficiencies by performing alignments of all samples to both reference genomes.

Supplemental Figure S4. Global read coverage distribution of cytosine in each context for all samples.

Supplemental Figure S5. Global distribution of methylation levels in each context for all samples.

Supplemental Figure S6. Global DNA methylation profiling at different genotype and stages of inflorescence.

Supplemental Figure S7. The expression levels of six gene groups for the methylation level analysis across gene regions.

Supplemental Figure S8. Influence of DNA methylation on gene expression at different genotype and stages of inflorescence.

Supplemental Figure S9. Differential DNA methylation regions between HAL2 and FIL2 inflorescences at D4 stage.

Supplemental Figure S10. Differential DNA methylation regions between D1 and D4 in HAL2 inflorescences.

Supplemental Figure S11. Differential DNA methylation regions between D1 and D4 in FIL2 inflorescences.

Supplemental Figure S12. Differential methylation and differential expression genes between HAL2 and FIL2 inflorescence.

Supplemental Figure S13. Association of differentially methylated genes with differentially expressed genes at D4 stage.

**Supplemental Table S1.** Factorial linear modeling analyses of genotype, development and their interaction for all expressed genes.

**Supplemental Table S2.** Significantly differentially expressed genes between HAL2 and FIL2 across four development stages and their clustering.

**Supplemental Table S3.** The quality and coverage of all bisulfite sequencing data.

**Supplemental Table S4.** Significantly CG-, CHG-, and CHH-differentially methylated genes between HAL2 and FIL2 at D1 stage.

**Supplemental Table S5.** Significantly CG-, CHG-, and CHH-differentially methylated genes between HAL2 and FIL2 at D4 stage.

**Supplemental Table S6.** Significantly CG-, CHG-, and CHH-differentially methylated genes between D1 and D4 stages in HAL2 inflorescence.

**Supplemental Table S7.** Significantly CG-, CHG-, and CHH-differentially methylated genes between D1 and D4 stages in FIL2 inflorescence.

## Acknowledgments

We thank the Department of Energy Joint Genome Institute for pre-publication access and permission to publish this analysis using the transcriptome and DNA methylome data of *Panicum hallii*. We thank Shane Merrell for growth chambers management.

